# Tracking the emergence of location-based spatial representations in human scene-selective cortex

**DOI:** 10.1101/547976

**Authors:** Sam C. Berens, Bárður H Joensen, Aidan J. Horner

## Abstract

Scene-selective regions of the human brain form allocentric representations of locations in our environment. These representations are independent of heading direction and allow us to know where we are regardless of our direction of travel. However, we know little about how these location-based representations are formed. Using fMRI representational similarity analysis and linear mixed-models, we tracked the emergence of location-based representations in scene-selective brain regions. We estimated patterns of activity for two distinct scenes, taken before and after participants learnt they were from the same location. During a learning phase, we presented participants with two types of panoramic videos: (1) an overlap video condition displaying two distinct scenes (0° and 180°) from the same location, and (2) a no-overlap video displaying two distinct scenes from different locations (that served as a control condition). In the parahippocampal cortex (PHC) and retrosplenial cortex (RSC), representations of scenes from the same location became more similar to each other only after they had been shown in the overlap condition, suggesting the emergence of viewpoint-independent location-based representations. Whereas these representations emerged in the PHC regardless of task performance, RSC representations only emerged for locations where participants could behaviourally identify the two scenes as belonging to the same location. The results suggest that we can track the emergence of location-based representations in the PHC and RSC in a single fMRI experiment. Further, they support computational models that propose the RSC plays a key role in transforming viewpoint-independent representations into behaviourally-relevant representations of specific viewpoints.

## Introduction

Rapidly learning the spatial layout of a new environment is a critical function that supports flexible navigation. This ability is thought to depend on the emergence of location-based representations in scene-selective brain regions that signal where we are irrespective of our current heading direction. As we are unable to sample all possible viewpoints from a given location simultaneously, the formation of location-based representations requires the integration of scenes from differing viewpoints. Despite evidence for the existence of location-based representations in scene-selective regions (e.g., Marchette, Vass, Ryan, & Epstein, 2015; Vass & Epstein, 2013), we know little about how such representations emerge.

Models of spatial navigation suggest that distinct brain regions are responsible for supporting allocentric (viewpoint-independent) and egocentric (viewpoint-dependent) representations of our environment (Byrne, Becker, & Burgess, 2007; Julian, Keinath, Marchette, & Epstein, 2018). Specifically, the parahippocampal cortex (PHC) and hippocampus are thought to encode allocentric spatial representations related to navigational landmarks/boundaries (Burgess, Becker, King, & O’Keefe, 2001; Epstein, Patai, Julian, & Spiers, 2017), and spatial context more broadly (Epstein & Vass, 2014). The hippocampus also supports a wider variety of spatial and non-spatial associative/configural functions in the service of memory and navigation (e.g. Eichenbaum, 2004; Hannula & Ranganath, 2009; Henson & Gagnepain, 2010; Kumaran et al., 2007; O’Keefe & Burgess, 2005). Here we focus on the parahippocampal cortex given its more specific role in spatial allocentric processing relative to the hippocampus. In contrast, the parietal lobe is thought to support egocentric representations of specific viewpoints that underpin route planning (Byrne et al., 2007; Calton & Taube, 2009). To enable efficient route planning, a transformation between allocentric and egocentric representations is thought to occur in the retrosplenial cortex, cueing allocentric representations from egocentric inputs and vice versa (Bicanski & Burgess, 2018; Byrne et al., 2007).

In support of these models, human fMRI studies using representational similarity analyses (RSA) have found evidence for viewpoint-independent representations of specific locations (henceforth referred to as “location-based representations”) in a network of brain regions including the PHC and RSC (Marchette, Vass, Ryan, & Epstein, 2014; Vass & Epstein, 2013). More recently, panoramic videos have been used to experimentally induce the formation of location-based representations (Robertson et al., 2016). Assessing pattern similarity for distinct scenes taken from the same location, Robertson et al. provided evidence for greater pattern similarity in the RSC and occipital place area (OPA) after participants had seen a panoramic video showing that two scenes were from the same location. This effect was not evident when participants could not learn that two scenes were from the same location. Interestingly, they also provided evidence for an effect in the PHC that occurred in both video conditions – i.e., regardless of whether participants could learn the scenes were from the same location – suggesting a more general associative role for the PHC.

Despite these results, we still know little about (*1*) how quickly such representations are formed, (*2*) what types of spatial information they encode, and (*8*) under what conditions they are evoked. First, it remains unclear whether location-based representations emerge rapidly after short exposures to a new environment, or whether they only develop after prolonged experience. Robertson et al. had participants watch videos outside of the scanner, over the course of two days, before assessing pattern similarity inside the scanner. To test whether location-based representations can form rapidly, we developed a protocol that permitted us to scan participants before and after a short learning phase, allowing us to estimate changes in pattern similarity as a function of learning in a single fMRI experiment. Second, without tracking the formation of location-based representations, it is difficult to determine exactly what type of information they are representing. For instance, shared representations across viewpoints may relate to long-term semantic knowledge that is invoked when seeing different views of a well-known location (see Marchette, Ryan, & Epstein, 2017). In contrast, rapidly learning representations that are shared across different viewpoints of a new environment implies that the information being encoded is more likely to be spatial rather than semantic in nature.

Third, we do not know whether location-based representations are involuntary retrieved during visual processing. Computational models of spatial navigation predict that allocentric representations are automatically activated and updated by egocentric viewpoints (Bicanski & Burgess, 2018; Byrne et al., 2007). Furthermore, electrophysiological studies in rodents have shown that allocentric representations are automatically activated and updated during exploration (e.g. Monaco, Rao, Roth, & Knierim, 2014; O’Keefe & Dostrovsky, 1971). However, evidence in humans is lacking. Robertson et al. required participants to recall whether scenes were presented on the left or right of the screen, introducing a task that explicitly required them to recall the panorama, and the position of the specific scene within the panorama. Suggesting some level of involuntary retrieval, one fMRI study found that viewpoint-independent representations of specific buildings may be activated when participants judge whether the building is well-known to them (Marchette et al., 2014). In the current study, participants performed an unrelated low-level attentional task as the scenes were presented. The activation of location-based representations under these conditions would suggest that they can be retrieved in a relatively automatic manner.

Here we test whether location-based representations of novel environments can be learnt by integrating visual information across different scenes. While location-based representations are predicted by models of spatial navigation, they may also be consistent with various other cognitive models (see Discussion). As such, we define location-based representations to be any type of information that encodes the relationship different, non-overlapping views of the same location. We recorded patterns of BOLD activity as participants passively observed a number of scenes depicting different views of novel locations. Subsequently, using an experimental manipulation introduced by Robertson et al. (2016), participants watched videos showing these scenes as part of a wider panorama. Half of the videos allowed participants to learn the spatial relationship between two scenes from the same location (overlap condition). The remaining videos acted as a control by presenting scenes from different locations (no-overlap condition). Following the videos, we again recorded patterns of activity for each of the scenes. Whereas Robertson et al. (2016) only assessed scene representations following video presentation, we also scanned before and during the videos; see Clarke, Pell, Ranganath, & Tyler (2016) for a similar pre- vs post-experimental design focussed on changes in object representations. This allowed us to track the potential *emergence* of location-based representations using representational similarity analyses (RSA), as well as assess neural activity when these representations were being formed.

Using generalised linear mixed models, we show that patterns evoked by different scenes become more similar in scene-selective regions of the PHC and RSC following the presentation of the video panoramas. This increase in similarity was specific to the ‘overlap’ video condition, where scenes from the same location were presented together, and was not observed in the no-overlap condition. This suggests the emergence of location-based representations in the PHC and RSC. Importantly, whereas this increase in pattern similarity emerged in the PHC regardless of behavioural performance, the same pattern was only present in the RSC when participants could remember which scenes came from the same location. This finding supports computational models that propose the RSC is critical in translating viewpoint-independent representations in the medial temporal lobe into more behaviourally-relevant egocentric representations.

## Methods

### Participants

Twenty-eight, right-handed participants were recruited from the University of York, UK. These participants had no prior familiarity with the locations used as stimuli in the experiment (see below). All participants gave written informed consent and were reimbursed for their time. Participants had either normal or corrected-to-normal vision and reported no history of neurological or psychiatric illness. Data from five participants could not be included in the final sample due to: problems with fMRI data acquisition (1 participant), excess of motion related artefacts in the imaging data (3 participants), and a failure to respond during one of the in-scanner tasks (1 participant). As such, analyses included 23 participants (10 males) with a mean age of 21.96 years (*SD* = 3.22). The study was approved by a local research ethics committee at the University of York.

### Stimuli

We generated 12 panoramic images of different urban locations from the City of Sunderland, and Middlesbrough town centre, UK (Figure 1; https://osf.io/cgy97). These panoramas spanned a 210° field-of-view horizontally but were restricted in the vertical direction to limit the appearance of proximal features (< 2 meters from camera). Throughout the experiment, 24 ‘endpoint images’ displaying 30° scenes taken from either end of each panorama were shown (i.e., centred at 0° and 180°; Figure 1A). These images were shown both inside and outside of the scanner to assess participants’ spatial knowledge of the depicted locations and for the representational similarity analysis (see below).

**Figure 1.**
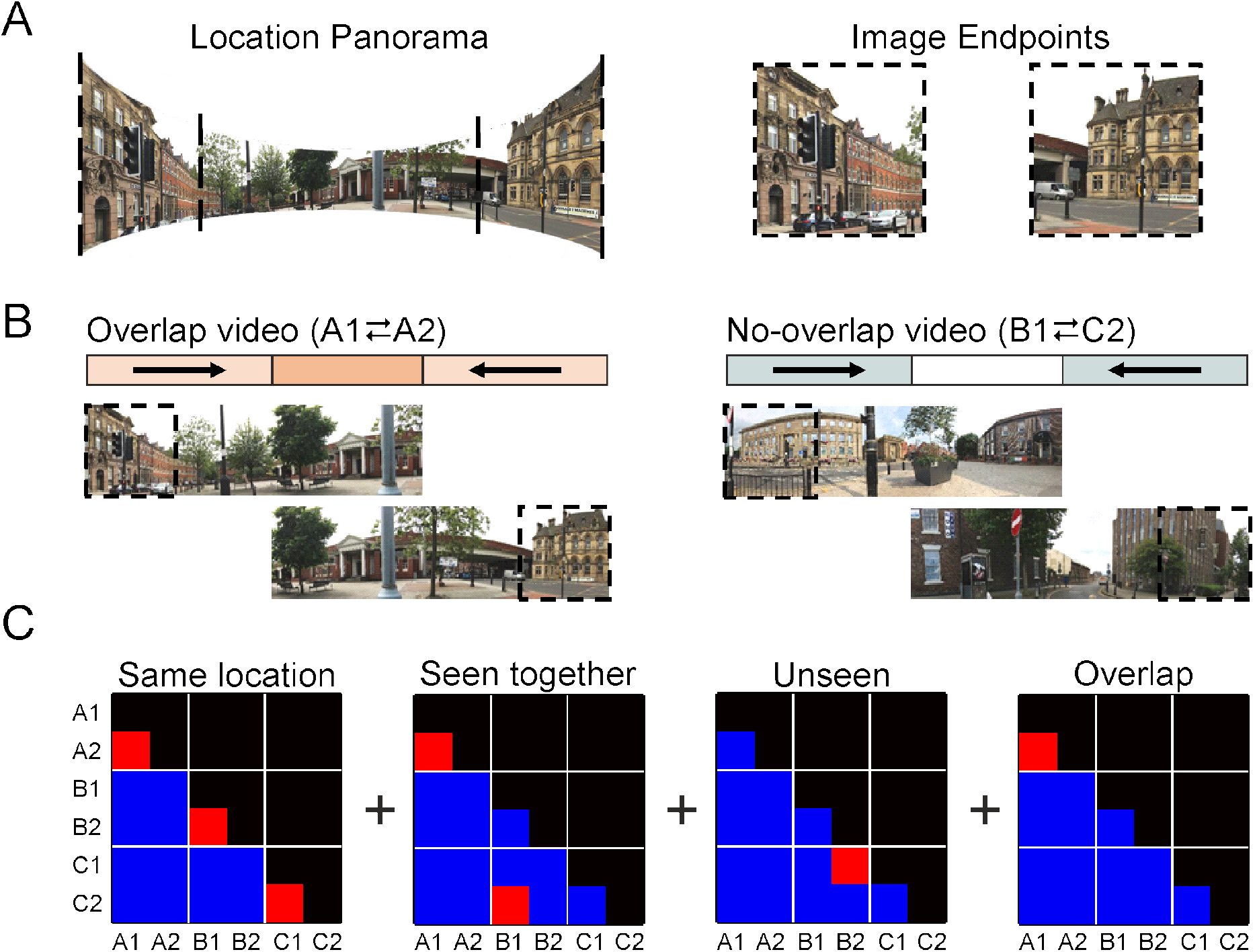
Stimuli used during, and analyses performed on, the in-scanner tasks. **(A)** An example location panorama with 2 endpoint images. Single endpoints were show during the in-scanner target detection task. As in Robertson et al. (2016), full panoramas were never shown as whole images but were presented during the in-scanner videos. **(B)** Depiction of the 2 video conditions: overlap vs no-overlap videos. Overlap videos showed camera pans from each endpoint of a given panorama (denoted A1 and A2) to the centre of that panorama. The central overlap allowed participants to learn a spatially coherent representation that included both A1 and A2. No-overlap videos involved pans between endpoints from two different locations (denoted B1 and C2) meaning that there was no visual overlap. **(C)** Similarity contrast matrices used to model changes in representational similarity between endpoints (i.e., between A1, A2, B1, B2, C1, and C2). Red squares indicate positively weighted correlations and blue squares indicate zero weighted correlations. From left to right, the matrices account for the representational similarity of endpoints: (1) from the same location regardless of video condition, (2) that were seen in the same video (including overlap and no-overlap videos), (3) in the unseen condition specifically, and (4) in the overlap condition specifically. Linear combinations of these matrices, along with their interactions with a session regressor (pre- vs post-videos), accounted for each RSA effect across all experimental conditions.

Endpoints were also shown in a series of videos (see https://osf.io/cgy97). In *overlap videos*, images A1 and A2 (taken from opposite ends of the same panorama) were presented such that their spatial relationship could be inferred (Figure 1B). Here, a camera panned from each endpoint to the centre of the panorama showing that A1 and A2 belonged to the same location. In contrast, a *no-overlap video* featured endpoints from two unrelated panoramas (images B1 and C2). Again, these videos showed an end-to-centre camera pan from each image. However, since there was no visual overlap between the video segments, observers could only infer that endpoints B1 and C2 belonged to different locations. The no-overlap condition acted as a control condition, ensuring endpoints B1 and C2 were seen in a similar video to endpoints in the overlap condition (A1 and A2), with the same overall exposure and temporal proximity. To ensure that the occurrence of a visual overlap was easily detectable, all videos alternated the end-to-centre sweep from each endpoint over two repetitions.

Pairs of endpoints from the same panorama were grouped into sets of 3. The first pair in each set were assigned to the overlap video condition (A1 and A2). Two endpoints from different panoramas were assigned to the no-overlap video condition (B1 and C2). The remaining endpoints belonged to an *unseen video* condition as they were not shown during any video (B2 and C1). These assignments were counterbalanced across participants such that each image appeared in all 3 conditions an equal number of times. The order of camera pans during videos (e.g., A1 first vs A2 first) was also counterbalanced both within and across participants. Analyses showed the visual similarity of image endpoints was matched across experimental conditions as measured by the Gist descriptor (Oliva & Torralba, 2001) and local correlations in luminance and colour information (https://osf.io/6sr9p/). Pilot data revealed that participants could not reliably identify which endpoints belonged to the same location without having seen the videos (https://osf.io/kgv64/).

### Procedure

Prior to entering the scanner, participants performed a behavioural task to assess their ability to infer which image endpoints were from the same location. Once in the scanner, they undertook a functional localiser task to identify scene-selective regions of the parahippocampal cortex (PHC) and retrosplenial cortex (RSC). They were then shown each image endpoint multiple times (performing a low-level attentional task) to assess baseline representational similarity between each image endpoint (i.e., prior to learning). During a learning phase, overlap and no-overlap videos were presented, with participants instructed to identify whether the endpoints in each video belonged to the same location or not. Following this video learning phase, each image endpoint was again presented multiple times to assess post-learning representational similarity between image endpoints. Finally, outside the scanner, participants performed the same behavioural task (as before scanning) to assess the extent to which participants had learnt which image endpoints belonged to the same location (and a further test of associative memory, see below). A figure illustrating the order and approximate duration of each experimental task is available at https://osf.io/zh8f2/.

### Pre-/Post-scanner tasks

Participants were tested on their ability to identify which endpoints belonged to the same location both before and after scanning (both outside of the scanner). On each trial, one endpoint surrounded by a red box was presented for 3 seconds. Following this, 5 other endpoints were displayed in a random sequence, each shown alongside a number denoting the order of appearance (i.e., 1-5; Figure 2A; 2 seconds per image, 500 ms inter-stimulus interval). One image in the sequence (the target) was taken from the same panorama as the cue. The remaining 4 endpoints (lures) belonged to panoramas in the same set of stimuli. As such, if endpoint B1 was presented as the cue, B2 would be the target, and A1, A2, C1, and C2 would be lures (i.e., a 5-alternative forced choice, 5-AFC). After the 5 alternatives had been shown, participants were prompted to select the target using a numeric key press (1-5). Across 24 trials, each endpoint was used as a cue image.

**Figure 2.**
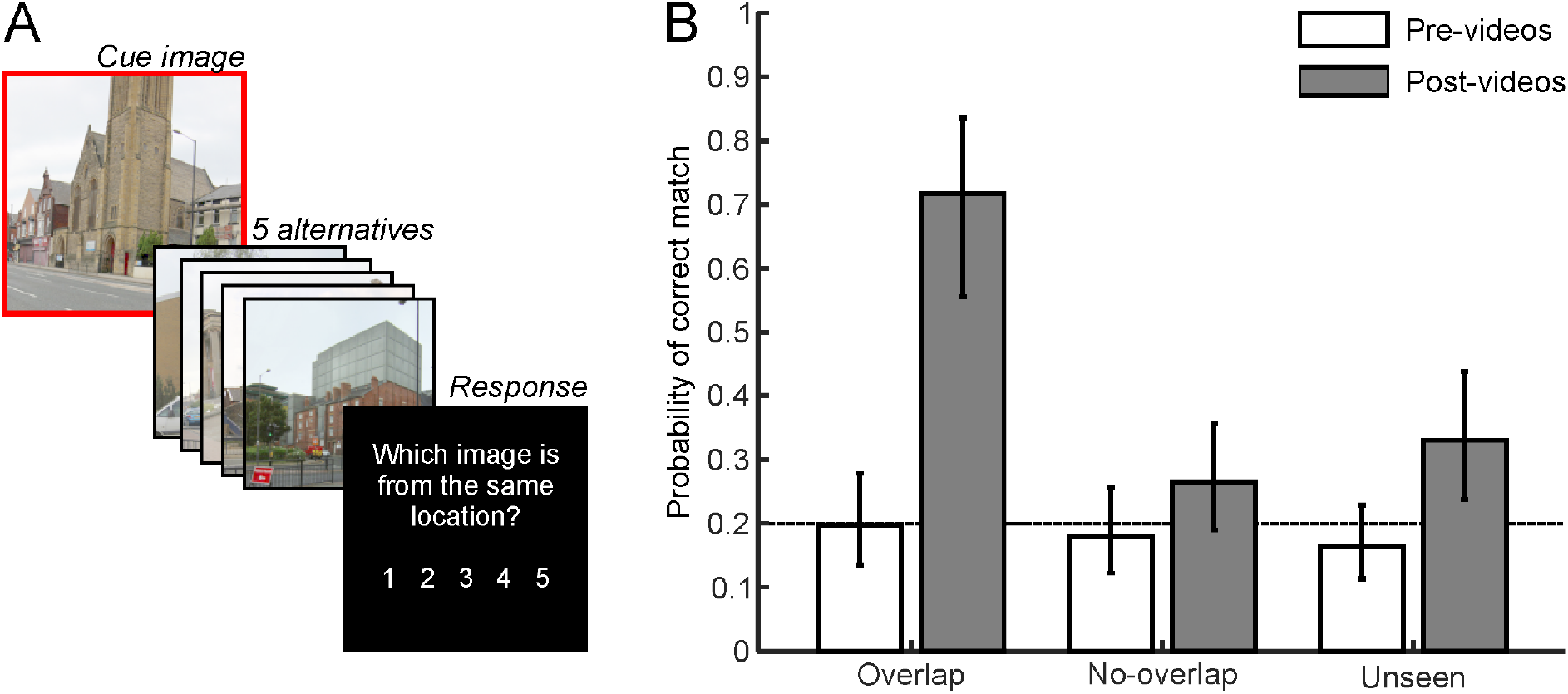
Behavioural task and results **(A)** A schematic illustration of the pre- and post-video behavioural task. One endpoint was first presented as a cue (enclosed by red box), followed by 5 numbered alternatives. Participants were then prompted to select which one of the alternatives belonged to the same location as the cue. **(B)** Performance on the pre- and post-video behavioural tasks plotted by video condition. Error bars represent 95% confidence intervals and the dashed line at *p* = .2 reflects 5-AFC chance level.

Following scanning, and the second block of the location identity task described above, participants were also asked to identify which images appeared together in the same video. Note that this is slightly different to the previous task since participants could have known that endpoints B1 and C2 appeared in the same video, despite not knowing which endpoints were from the same location (i.e., B2 and C1 respectively). Using a similar procedure to that described above, endpoints from either the overlap or no-overlap video conditions were cued and participants were asked to select the appropriate endpoint from the 5 alternatives in the same set.

### In-scanner tasks

#### Functional localiser

Before the main experimental task, participants undertook a functional localiser scan with the purpose of identifying four scene selective regions of interest (ROIs) - in particular, the left and right parahippocampal cortex (PHC), and the left and right retrosplenial cortex (RSC). This involved presenting 4 blocks of scene images (coasts, mountains, streets, and woodlands) interleaved with 4 blocks of face images (male and female). In each block, 10 unique images were shown in quick succession with a display time of 700 ms per image and an inter-stimulus interval of 200 ms. Blocks were separated with a 9 second inter-block interval and their running order was counterbalanced across participants. The scene images used here were different to those in the main experiment and none were repeated during the localiser itself. All images were shown in greyscale and were presented with a visual angle of ~14°. To ensure localiser images were being attended to, participants were tasked with detecting an odd-ball target that was superimposed onto one of the images in each block. The target was a small red dot with a 3-pixel radius. When this was seen, participants were required to respond with a simple button press as quickly as possible (mean detection performance: *d’* = 3.116, *SD* = 0.907).

#### Presentation of endpoint images

Participants were shown all 24 endpoint images during an event-related functional imaging task. The task was optimised to measure multivariate patterns of BOLD activity specific to individual endpoints and was run both before and after participants had seen the panoramic videos (session 1: pre-videos; session 2: post-videos). All endpoints were presented 9 times for both the pre- and post-video functional run. Images were displayed for 2.5 seconds with an inter-stimulus interval of 2 seconds. The order of stimuli in each functional run was optimised to facilitate the decoding of unique BOLD patterns across endpoints (optimisation algorithm available at https://osf.io/eh78w/). No image was presented on successive trials to avoid large adaptation effects and the design included 12 null events in each functional run (i.e., 10% of all events). Like the functional localiser, participants were tasked with detecting an odd-ball target that was superimposed onto a small proportion of the images. Here, the target was a group of 3 small red dots (3-pixel radius, < 0.2°), with each dot drawn at a random position on the image. Targets were present on 1 out of every 9 trials such that 8 repetitions of each endpoint image were target free (target trials were not used to estimate BOLD patterns). As above, participants were required to respond to these targets with a simple button press (mean detection performance, *d’*, was 3.362, *SD* = 0.642, pre-videos, and 3.659, *SD* = 0.485, post-videos).

#### Panoramic video task

Participants watched all video clips from the overlap and no-overlap video conditions whilst being scanned. Each video lasted a total of 20 seconds and was followed by a 10 second rest period. In the first 3 seconds of this rest period, participants were prompted to indicate whether each video segment depicted scenes from the same or different locations. Responses were recorded with a left/right button press. This question was asked to ensure that participants were attending to the visual overlap across segments (mean discrimination performance: *d’* = 3.220, *SD* = 0.373). All videos were repeated 3 times in a pseudorandom order to allow for sufficient learning. Prior to entering the scanner, participants were asked to remember which endpoints were seen together in the same video, even if they appeared in a no-overlap video. Participants were told that a test following the scan would assess their knowledge of this.

### MRI acquisition

All functional and structural volumes were acquired on a 3 Tesla Siemens MAGNETOM Prisma scanner equipped with a 64-channel phased array head coil. T2*-weighted scans were acquired with echo-planar imaging (EPI), 35 axial slices (approximately 0° to AC-PC line; interleaved) and the following parameters; repetition time = 2000 ms, echo time = 30 ms, flip angle = 80°, slice thickness = 3 mm, inplane resolution = 3 × 3 mm. The number of volumes acquired during (a) the functional localiser, (b) the video task, and (c) each run of the endpoint presentation task was 75, 363, and 274, respectively. To allow for T1 equilibrium, the first 3 EPI volumes were acquired prior to the task starting and then discarded. Subsequently, a field map was captured to allow the correction of geometric distortions caused by field inhomogeneity (see the MRI pre-processing section below). Finally, for purposes of co-registration and image normalization, a whole-brain T1-weighted structural scan was acquired with a 1mm^3^ resolution using a magnetization-prepared rapid gradient echo pulse sequence.

### MRI pre-processing

Image pre-processing was performed in SPM12 (www.fil.ion.ucl.ac.uk/spm). This involved spatially realigning all EPI volumes to the first image in the time series. At the same time, images were corrected for field inhomogeneity based geometric distortions (as well as the interaction between motion and such distortions) using the Realign and Unwarp algorithms in SPM (Andersson, Hutton, Ashburner, Turner, & Friston, 2001; Hutton et al., 2002). For the RSA, multivariate BOLD patterns of interest were taken as *t*-statistics from a first-level general linear model (GLM) of unsmoothed EPI data in native space. Aside from regressors of interest, each first-level GLM included a set of nuisance regressors: 6 affine motion parameters, their first-order derivatives, regressors censoring periods of excessive motion (rotations > 1°, and translations > 1mm), and a Fourier basis set implementing a 1/128 Hz high-pass filter. For the analyses of univariate BOLD activations, EPI data were warped to MNI space with transformation parameters derived from structural scans (using the DARTEL toolbox; Ashburner, 2007). Subsequently, the EPI data were spatially smoothed with an isotropic 8 mm FWHM Gaussian kernel prior to GLM analysis (regressors included the same nuisance effects noted above).

### Regions of interest

We generated four binary masks per participant to represent each ROI in native space. To do this, a first-level GLM of the functional localiser data modelled BOLD responses to scene and face stimuli presented during the localiser task. Each ROI was then defined as the conjunction between a “scene > face” contrast and an anatomical mask of each region that had been warped to native space (left/right PHC sourced from: Tzourio-Mazoyer et al., 2002; left/right RSC sourced from: Julian, Fedorenko, Webster, & Kanwisher, 2012). Thus, the ROIs were functionally defined but constrained to anatomical regions known to be spatially selective. Normalised group averages of these ROIs are available at https://osf.io/gbznp/ and https://neurovault.org/collections/4819.

Recent evidence suggests that the RSC is composed of at least two functionally distinct sub-regions, both of which may be scene-selective: (1) a retinotopically organised medial-place area in posterior sections of the retrosplenial complex, and (2) a more anterior region corresponding to BA29 and BA30 associated with more integrative mnemonic processes (Silson, Steel, & Baker, 2016). In the current study, we focus on the functionally defined retrosplenial cortex as a whole and do not differentiate between these sub-regions. However, the functional ROIs that we identified for each participant principally cover anterior sections of the RSC corresponding to BA29 and BA30, and show little overlap with the retinotopic areas identified by Silson et al.

The occipital place area (OPA) has also been implicated as a critical scene-selective region (e.g., Marchette et al., 2015; Robertson et al., 2016). Recent research suggests that this region is principally involved in representing environmental boundaries and navigable paths during visual perception (Bonner & Epstein, 2017; Julian, Ryan, Hamilton, & Epstein, 2016; Malcolm, Silson, Henry, & Baker, 2018). However, computational models of spatial navigation do not predict that the OPA maintains location-based representations that are viewpoint-invariant (Bicanski & Burgess, 2018; Byrne et al., 2007). Additionally, we were only able to reliably delineate the OPA bilaterally in 6 out of the 23 participants in our sample. As such, we did not focus on this region in the current study; instead, we restricted our main analyses and family-wise error corrections to the PHC and RSC bilaterally. Nonetheless, for completeness, we generated an OPA mask using a normalised group-level contrast and ran the location-based RSA analyses reported below on this region separately (statistical outputs available at https://osf.io/d8ucj/). No effects of interest were identified in either the left or right OPA.

### Representational similarity analyses

Our general approach to the RSA involved modelling the observed similarity between different BOLD patterns as a linear combination of effects of interest and nuisance variables. Here, the similarity between BOLD responses was taken as the correlation of normalised voxel intensities (*t-*statistics) across all voxels in an ROI. The resulting correlation coefficients were then Fisher-transformed before being subjected to statistical analysis. This transform ensures that the sampling distribution of similarity scores is approximately normal in order to meet the assumption of normality for statistical inference. We then entered all the transformed similarity scores under test from each participant and stimulus set into a general linear mixed-effects regression model. Whilst underused in the neurosciences (though see Motley et al., 2018), these models are common in the psychological literature as they offer a robust method of modelling non-independent observations with few statistical assumptions (Baayen, Davidson, & Bates, 2008). Here, we used mixed-effects models to predict observed representational similarity between endpoints with a set of fixed-effects and random-effects predictors (discussed below).

Importantly, mixed-effects models allow us to include estimates of pattern similarity across individual items (endpoints) and participants in the same statistical model. The fixed-effects predictors in each model specified key hypotheses of interest. The random effects accounted for statistical dependencies between related observations at both the item- and participant-level. RSA analyses of fMRI data typically either assess patterns across all items (regardless of condition), or average across items in the same condition, meaning that important variation within conditions is ignored. Our modelling approach allows us to examine changes in representational similarity at the level of both items and conditions simultaneously while controlling for statistical dependencies between related observations.

Raw similarity data and mean similarity matrices are available on the Open Science Framework (OSF; https://osf.io/cgy97). This page also includes MATLAB functions for estimating each statistical model as well as the model outputs.

### Visual representations of specific endpoints

We first examined whether the passive viewing of endpoint images evoked stimulus specific visual representations in each of our four ROIs (left and right PHC and RSC). Multivariate BOLD responses to the endpoints were estimated for session 1 (pre-videos) and session 2 (post-videos) separately. We then computed the similarity of these responses across sessions by correlating BOLD patterns in session 1 with patterns in session 2. This resulted in a non-symmetric, 24 x 24 correlation matrix representing the similarity between all BOLD patterns observed in session 1 and those observed in session 2. The correlation coefficients (n = 576 per participant) were then Fisher-transformed and entered as a dependent variable into a mixed-effects regression model with random effects for participants and endpoints. The main predictor of interest was a fixed effect that contrasted correlations between the same endpoints (e.g. A1-A1, B1-B1, n = 24 per participant) with correlations between different endpoints (e.g. A1-A2, A1-B1 etc., n = 552 per participant) across the two sessions.

As well as running this analysis in each ROI, we performed a complementary searchlight analysis to detect endpoint-specific representations in other brain regions. Here, local pattern similarity was computed for each brain voxel using spherical searchlights with a 3-voxel radius (the mean number of voxels per searchlight was 105.56; searchlights were not masked by grey-/white-matter tissue probability maps). Fisher-transformed correlations for same-versus different-endpoints were contrasted at the first-level before running a group-level random effects analysis.

### Location-based memory representations

We next tested our principal hypothesis – whether representations of endpoints A1 and A2 became more similar to one another as a result of watching the overlap videos – in each ROI. Using the multivariate BOLD responses from sessions 1 and 2, we computed the neural similarity between endpoints that were presented within the same image set and the same session. This resulted in 8 symmetric, 6 x 6 correlation matrices for each participant – one per set in session 1, and one per set in session 2. All the correlation coefficients from the lower triangle of these matrices (n = 15) were then Fisher-transformed and entered as a dependent variable into a mixed-effects regression model (see Figure 1C). As such, the model included a total of 120 correlations coefficients per participant (2 sessions x 4 sets x 15 similarity scores).

One fixed-effects predictor modelled unspecific changes in similarity between sessions (hereafter referred to the session effect) by coding whether similarity scores were recorded in session 1 or session 2. Similarly, a further 3 fixed-effects predictors modelled similarity differences attributable to: (*1*) endpoints in the overlap condition (i.e. A1-A2), (*2*) endpoints shown in the same video (A1-A2, B1-C2), and (*3*) endpoints that were not shown in any video (C1-B2) – shown in Figure 1C. Together, these predictors and their interactions constituted a 2×3 factorial structure (session [1 vs 2] by condition [overlap vs no-overlap vs unseen]) and so were tested with a *Session*Condition F-*test. Nonetheless, our principal hypothesis holds that there will be a specific interaction between the *Session* and *Overlap* predictors (referred to as the *Session*Overlap* effect) which we report alongside the *F*-test. The model also included a predictor indicating whether endpoints were from the same location (A1-A2, B1-B2, C1-C2) thereby allowing us to estimate changes in similarity between them. This ensured that variance loading onto the *Session*Overlap* effect was properly attributable to the learning of spatially coherent representations rather than some combination of other factors (e.g. same location + seen in same video). Note, this model term quantifies similarity differences between overlap endpoints and all other endpoints that *change* between session 1 and session 2. A positive effect may indicate either an increase in similarity in the overlap condition, or a decrease across all other similarity scores regardless of condition (or both). As such, the model is structured to account for any systematic change in the baseline level of similarity across sessions (see results). Furthermore, the *Session*Overlap* term is only sensitive to a learning effect that causes relative shifts in similarity scores specific to the overlap condition and cannot be attributed to any other combination of effects.

Finally, the model included a behavioural predictor specifying whether participants were able to match endpoints A1 to A2 in the post-scanner task (mean centred with 3-levels: 0, 1 or 2 correct responses per pair). This examined whether changes in representational similarity were dependent on participants’ ability to identify that endpoints from the overlap condition belonged to the same location after scanning (i.e., a 3-way interaction; *Session*Overlap*Behaviour*). Random effects in the model accounted for statistical dependencies across image sets, sessions, and subjects.

To complement the ROI analyses, we ran a searchlight analysis that tested for RSA effects across the whole brain (searchlight radius: 3-voxel). Here, first-level contrast estimates compared the Fisher-transformed correlations between overlap endpoints (i.e. A1-A2) and all other endpoint correlations (e.g. B1-B2, B1-C1). A group-level analysis then compared these similarity contrasts between sessions to test the *Session*Overlap* interaction. To test for a *Session*Overlap*Behaviour* interaction, the group-level model also included a behavioural predictor specifying participant’s average performance in matching A1 to A2 during the post-scanner task (mean centred). Note that this searchlight analysis is not able to control for the potential contributions of other important factors (i.e., same-location, same-video) that our mixed-effects approach explicitly controls. It is complementary, but secondary, to the ROI analyses.

### Statistical validation and inference

To ensure that each mixed-effects regression model was not unduly influenced by outlying data points, we systematically excluded observations that produced unexpectedly large residuals more than 2.5 standard deviations above or below model estimates. This was conducted regardless of condition, so did not bias the analyses to finding an effect (if no effect were present). Further, a highly similar pattern of results was seen when not excluding outliers, supporting the robustness of our findings (see https://osf.io/dzy3p). Following these exclusions, Kolmogorov–Smirnov tests indicated that residuals were normally distributed across all the linear mixed-effects models. Additionally, visual inspection of scatter plots showing residual versus predicted scores indicated no evidence of heteroscedasticity or non-linearity. Where effects size estimates are contrasted across different models, we report the result as an unequal variances *t*-test with the degrees of freedom being approximated using the Welch–Satterthwaite equation (Welch, 1947).

All *p*-values are reported as two-tailed statistics. Family-wise error corrections related to the multiple comparisons across our 4 regions of interest are made for each *a priori* hypothesis (denoted *p_FWE_*). Additionally, we report whole-brain effects from searchlight and mass univariate analyses when they survive family-wise error corrected thresholds (*p_FWE_* < .05) at the cluster level (cluster defining threshold: *p* < .001 uncorrected). All other *p*-values are noted at uncorrected levels. As well as reporting null hypothesis significance tests, we present the results of complimentary Bayesian analyses. Unlike the frequentist statistics, these indicate whether the null is statistically preferred over the alternative hypothesis. As such, we use the Bayesian analyses to determine whether there is evidence for the null when frequentist tests are non-significant. For each *t*-test, a Bayes factor in favour of the null hypothesis (*BF_01_*) was computed with a Cauchy prior centred at zero (i.e., no effect) and a scale parameter (*r*) of 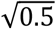 (see Gelman, Jakulin, Pittau, & Su, 2008). Bayes factors greater than 3 are taken as evidence in favour of the null hypothesis while those less than 1/3 are taken as evidence in favour of the alternative (Kass & Raftery, 1995). Finally, alongside the inferential statistics, we report Cohen’s *d* effect sizes for each *t*-test. When effects are tested in the context of a mixed-effects model, estimates of Cohen’s *d* are computed from the fixed-effects only and exclude variance attributed to random-effects.

## Results

### Behavioural performance

We first analysed behavioural responses to the pre- and post-scanner tasks to determine (a) whether participants were able to identify which endpoints belonged to the same location, and (b) whether performance increased as a result of watching the overlap videos. A generalised linear mixed-effects analysis modelled correct vs incorrect matches between cue and target endpoints as a function of session (pre- vs post-videos) and experimental condition (overlap, no-overlap, and unseen). As such, the model constituted a 2 x 3 factorial design with random intercepts and slopes for both participants and endpoints.

The results, displayed in Figure 2B, revealed significant main effects of session (*F_1, 1098_* = 47.302, *p* < .001), and condition (*F_2,1098_* = 6.500, *p* = .002), as well as an interaction between them (*F_2, 1098_* = 11.231, *p* < .001). The interaction indicated that performance was at chance level across all conditions before the videos (min *p* = 0.249, *BF_01_* = 2.531, *d* = −0.241), but substantially increased in the overlap video condition after the videos (*t_1098_* = 6.867, *p* < .001, *BF_01_* < 0.001, *d* = 1.432; post-video > pre-video). This increase was not seen in the no-overlap condition (*t_1098_* = 1.761, *p* = .079, *BF_01_* = 1.212, *d* = 0.3672), however a significant increase was seen in the unseen condition (*t_1098_* = 3.159, *p* = .002, *BF_01_* = 0.105, *d* = 0.659). The performance increases in the control conditions (only significant in the unseen condition) were likely the result of participants being able to exclude overlap endpoints as non-target alternatives in the 5-AFC test (i.e., a recall-to-reject strategy, disregarding A1 and A2 when cued with either B1, B2, C1 or C2). Consistent with this, session 2 performance in the no-overlap and unseen conditions was not significantly different from chance level in a 3-AFC test (0.33; as opposed to 0.2 in a 5-AFC test; no-overlap: *t_1098_* = −1.494, *p* = .135, *BF_01_* = 1.729, *d* = −0.312; unseen: *t_1098_* = −0.054, *p* = .957, *BF_01_* = 4.567, *d* = −0.011). Nonetheless, performance in the overlap condition did significantly differ from this adjusted chance level (*t_1098_* = 4.514, *p* < .001, *BF_01_* = 0.006, *d* = 0.941).

Participants’ increased ability to match endpoints in the overlap condition was not characteristic of a general tendency to match endpoints that appeared in the same video (i.e., selecting B1 when cued with C2). This was evident since matches between no-overlap endpoints were not more likely in session 2 compared with session 1 (*t_366_* = 0.646, *p* = .519, *BF_01_* = 3.785, *d* = 0.135). In contrast, performance increases in the overlap condition (i.e., the post-video > pre-video effect reported above) were significantly larger than this general effect of matching all endpoints that appeared in the same video (*t_949,20_* = 5.027, *p* < .001, *BF_01_* = 0.002, *d* = 1.048). Additionally, participants were unable to explicitly match no-overlap endpoints shown in the same video during the final behavioural task (comparison to 0.2 chance level: *t_334_* = −0.467, *p* = .641, *BF_01_* = 4.141, *d* = −0.097). In sum, participants rapidly learnt which scenes were from the same location, however this was only seen in the overlap condition (and not in the no-overlap condition).

### Visual representations of specific endpoints

First, we report the results of the mixed-effects model testing for representations of specific endpoints that remained relatively unchanged across sessions (i.e., pre- to post-videos). This revealed that correlations between the same endpoints (e.g., A1-A1, B1-B1) were greater than correlations between different endpoints (e.g., A1-A2, A1-B1) in both the right PHC (*t_13224_* = 5.229, *p_FWE_* < .001, *BF01* = 0.001, *d* = 1.090), and the left PHC (*t_13200_* = 6.351, *p_FWE_* < .001, *BF_01_* < 0.001, *d* = 1.324). This effect was not significant in either the right or left RSC (*t_13210_* = 1.185, *p_FWE_* = .945, *BF_01_* = 2.454, *d* = 0.247, and *t_13202_* = −0.231, *p_FWE_* = .999, *BF_01_* = 4.463, *d* = −0.048, respectively).

The searchlight analysis that tested for consistent representations of specific endpoints across the whole brain revealed representations in one large cluster that peaked in the right occipital lobe (area V1; *t_22_* = 11.50, *p_FWE_* < .001, *k* = 5202, *BF_01_* < 0.001, *d* = 2.398) and extended into the areas V2, V3, V4, and the fusiform gyri bilaterally. Three smaller clusters were also detected in the right precuneus (*t_22_* = 4.64, *p_FWE_* = .011, *k* = 44, *BF_01_* = 0.005, *d* = 0.968), right inferior parietal lobule (*t_22_* = 4.40, *p_FWE_* = .028, *k* = 37, *BF_01_* = 0.008, *d* = 0.918), and right RSC (*t_22_* = 4.32, *p_FWE_* = .025, *k* = 38, *BF_01_* = 0.008, *d* = 0.901). The latter effect overlapped considerably with the right RSC region of interest identified for each subject. However, the effect size estimated in the ROI analysis was weaker than the peak searchlight effect, principally because it was variable across endpoints and as such largely accounted for by random effects in the model. Unthresholded statistical maps of these effects are available at https://neurovault.org/collections/4819.

In sum, we find evidence that the PHC (bilaterally), the right RSC, and a number of early visual areas maintained consistent representations of specific endpoints across scanning sessions. Note, whether a region codes such representations across scanning sessions is independent of whether it may learn location-based memory representations in the second session; these effects are, in principle, dissociable.

As part of a supplementary analysis, we also tested for visual representations of specific scenes that remined stable within (but not necessarily across) scanning sessions (see https://osf.io/exzba/). To quantify the BOLD similarity of specific scenes within each session, we required 2 independent pattern representations per session. Thus, across both sessions, we estimated voxel patters derived from 4 distinct periods: *a*) 1^st^ half of session 1 (pre-videos), *b*) 2^nd^ half of session 1 (pre-videos), *c*) 1^st^ half of session 2 (post-videos), and *d*) 2^nd^ half of session 2 (post-videos). As a result, each of these voxel representations was only derived from 4 endpoint presentations. Nonetheless, when similarity scores were modelled in a mixed-effects regression, each ROI showed greater levels of similarity between representations of the same endpoint relative to the similarity between different endpoints (weakest effect in the left PHC: *t_26492_* = 2.211, *p* = .027, *BF_01_* = 0.606, *d* = 0.461). Furthermore, this analysis revealed that representations of the same endpoints became more similar to one other after the videos in the right RSC and left PHC (weakest effect: *t_26492_* = 2.598, *p* = .009, *BF_01_* = 0.308, *d* = 0.542). This latter effect was insensitive in the right PHC and left RSC (weakest effect: *t_26492_* = 1.671, *p* = .095, *BF_01_* = 1.375, *d* = 0.348).

### Location-based memory representations

#### Effects in the right PHC

Next, we report the results of the mixed-effects model examining whether pattern similarity between different endpoints changed across sessions as a result of watching the videos. This revealed a significant *Session*Condition* interaction in the right PHC (*F_2, 2739_* = 6.827, *p_FWE_* = .004; average similarity matrices shown in Figure 3; condition estimates and confidence intervals plotted in Figure 4A). Post-hoc tests showed that this effect was driven by a difference between pre- to post-video sessions for endpoints in the overlap condition (*t_2739_* = 2.923, *p* = .004, *BF_01_* = 0.167, *d* = 0.610). This difference was not observed in any other condition (no-overlap: *t_2739_* = 0.756, *p* = .450, *BF_01_* = 3.533, *d* = 0.156; unseen: *t_2739_* = −0.970, *p* = .332, *BF_01_* = 3.001, *d* = −0.202). Furthermore, a significant *Session*Overlap* interaction highlighted that the similarity differences in the overlap condition was attributable to the video manipulation alone rather than some combination of other factors (*t_2739_* = 2.549, *p_FWE_* = .043, *BF_01_* = 0.337, *d* = 0.532).

**Figure 3.**
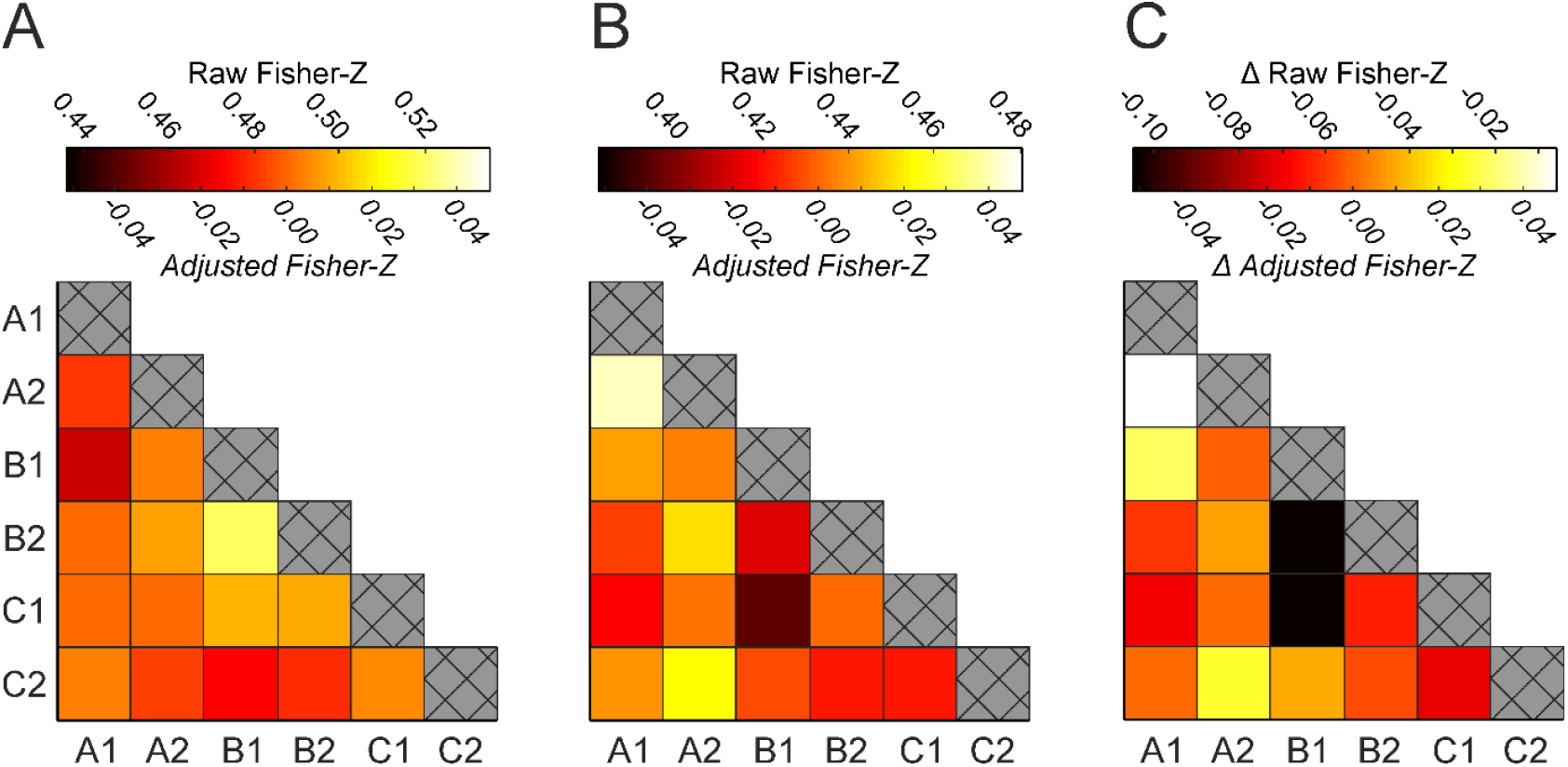
Mean representational similarity between endpoints in the right PHC, averaged across all participants and image sets. **(A)** Similarity between endpoints *before* the panoramic videos were shown (i.e. in session 1). **(B)** Similarity between endpoints *after* the panoramic videos were shown (i.e. in session 2). **(C)** Change in similarity that followed the panoramic videos (i.e. session 2 minus session 1). Colour bars indicate both raw and baseline-adjusted Fisher-Z statistics (above and below the colour bar respectively). Adjusted statistics account for trivial differences in similarity across scanning sessions caused by motion and scanner drift. This is achieved by subtracting out a baseline level of similarity between non-associated endpoints (i.e. endpoints that were not from the same location, video, or experimental condition). Note, the baseline-adjusted statistics are shown for illustrative purposes only; each RSA was conducted on the raw Fisher-Z statistics alone. Crosshatchings along the diagonal elements represent perfect correlations between identical BOLD responses and so were not included in the analyses.

**Figure 4.**
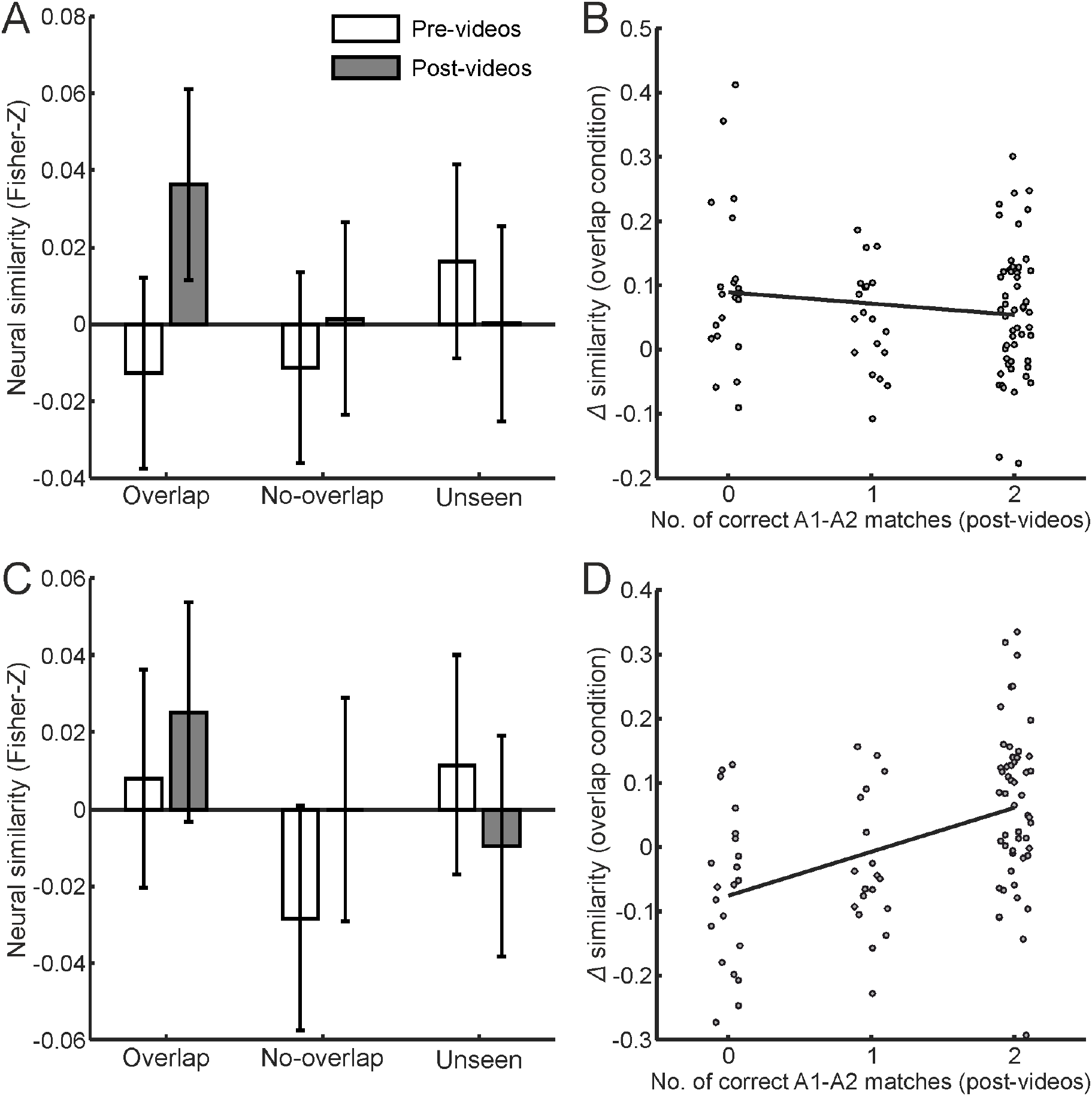
Results of the representational similarity analyses in the right parahippocampal cortex (rPHC, top row) and right retrosplenial cortex (rRSC, bottom row). **A:** rPHC similarity estimates of scenes in the pre- and post-video sessions, plotted by experimental condition. There was a significant change in similarity estimates between sessions in the overlap condition (*t_2739_* = 2.923, *p* = .004, *BF_01_* = 0.167, *d* = 0.610) that was not present in the no-overlap and unseen conditions (*t_2739_* = 0.756, *p* = .450, *BF_01_* = 3.533, *d* = 0.158, and *t_2739_* = −0.970, *p* = .332, *BF_01_* = 3.001, *d* = −0.202 respectively). **B:** In the rPHC, pre- to post-video changes in representational similarity for the overlap condition plotted against the number of correct matches between overlap endpoints in the post-video behavioural task. This association was not significant (*t_2739_* = −0.892, *p* = .373, *BF_01_* = 3.199, *d* = −0.186). **C & D:** Same as panels A and B but for the rRSC region of interest. The rRSC showed no overall similarity changes in any of the experimental conditions (*t_2728_* = 0.870, *t_2728_* = 1.419, and *t_2728_* = −1.059 for the overlap, no-overlap and unseen conditions respectively, all *p’s* > .156, *BF_01_’s* > 1.895, *d’s* < 0.296). Nonetheless, there was a significant association between behavioural performance and similarity changes in the overlap condition (*t_2728_* = 2.886, *p* = .004, *BF_01_* = 0.179, *d* = 0.602). All bars plot baseline corrected similarity estimates having subtracted out correlations between non-associated endpoints (e.g., A1-B1, A1-B2, etc.). As such, the zero line in panels A and C denotes the average similarity of these non-associated endpoints in each session. Error bars indicate 95% confidence intervals.

Importantly, before the videos were shown, pairs of endpoints from the same location (i.e., A1-A2, B1-B2 and C1-C2) were found to evoke neural patterns that were more similar to each other than pairs of endpoints from different locations in the right PHC (e.g., A1-B2, B1-C2; *t_2739_* = 2.498, *p_FWE_* = .050, *BF_01_* = 0.369, *d* = 0.521, see https://osf.io/uxhs9 for a plot of this effect). This ‘same-location’ effect suggests that even before the spatial relationship between scenes were known, the right PHC encoded visual properties of those scenes that generalised across different views. These data demonstrate that despite controlling for similarity across stimuli using both the GIST descriptor and a pixel-wise correlation, and despite participants being unable to infer which endpoints were from the same location prior to watching the videos, we still found evidence for a ‘same-location’ effect in the right PHC. This underlies the critical role of estimating pattern similarity prior to learning to identify significant increases in similarity post-video relative to pre-video (c.f., Robertson et al., 2016). Note, this ‘same-location’ effect is only seen when collapsing across all endpoint pairs, and is not evident in the session 1 *Overlap* condition alone (https://osf.io/uxhs9).

#### Effects in the right RSC

The *Session*Condition* and *Session*Overlap* interactions were not significant in any other ROI (*F’s* < 2.775, *p_FWE_’s* > .250; similarity estimates for the right RSC plotted in Figure 4C). However, we saw a significant *Session*Overlap*Behaviour* interaction in the right RSC (*t_2728_* = 2.886, *p_FWE_* = .016, *BF_01_* = 0.179, *d* = 0.602; Figure 4D). This suggests that the RSC only encoded viewpoint-independent representations when the spatial relationships between endpoints could be retrieved during the post-scanner test. No other ROIs showed a significant *Session*Overlap*Behaviour* interaction (largest effect: *t* = 0.050, *p_FWE_* = 1, *BF_01_’s* = 4.567, *d* = −0.010).

#### Differentiating the PHC and RSC

We next assessed whether there was evidence for dissociable roles of the right PHC and RSC, given that both represented location-based information but were differently associated with behavioural performance. Specifically, we assessed whether location-based representations in the RSC were significantly more associated with participants’ ability to match endpoints from the same location compared to representations in the PHC. This would suggest that the RSC plays a greater role in guiding behavioural performance than the PHC. We therefore tested whether the *Session*Overlap*Behaviour* (3-way) effect was larger in the RSC than the PHC. A comparison of effect sizes did show evidence for such a dissociation (*t_5311.9_* = 3.931, *p* < .001, *BF_01_* = 0.021, *d* = 0.820).

This implies that the right PHC might have exhibited above-baseline pattern similarity between A1 and A2 endpoints even when those endpoints were not subsequently remembered as belonging to the same location. We directly tested this by re-running the RSA having excluded A1/A2 pairs that were consistently remembered as belonging to the same location (i.e., having 2 correct responses during the post-scanner test). Despite these exclusions, pattern similarity differences in the overlap condition remained significant (*t_1188_* = 2.364, *p* = .018, *BF_01_* = 0.528, *d* = 0.493) and were not seen in any other condition (no-overlap: *t_1188_* = 0.324, *p* = .746, *BF_01_* = 4.359, *d* = 0.068; unseen: *t_1188_* = −0.585, *p* = .559, *BF_01_* = 3.915, d = −0.122); see Figure 4B which plots the size of the *Session*Overlap* effect in the right PHC at each level of behavioural performance. In contrast, the right RSC only showed above-baseline pattern similarity when the endpoints were consistently remembered as belonging to the same location. Re-running the RSA on these remembered pairs alone revealed similarity increases between consistently remembered endpoints in the overlap condition (*t_1538_* = 2.449, *p* = .014, *BF_01_* = 0.402, *d* = 0.511, see Figure 4D) that were not seen in any other condition (no-overlap: *t_1538_* = 1.107, *p* = .269, *BF_01_* = 2.651, *d* = 0.230; unseen: *t_1538_* = −1.316, *p* = .188, *BF_01_* = 2.134, *d* = −0.274).

In sum, we saw an increase in pattern similarity in the right PHC and right RSC between different scenes of the same location after they had been presented in an overlap video. Further, we observed a dissociation between the PHC and the RSC. Whereas the PHC showed increased pattern similarity regardless of performance on the post-scanner test, the RSC only showed increased pattern similarity when participants were able to subsequently identify those scenes as belonging to the same location.

#### Across session decreases in pattern similarity

Our mixed-effects regression models were conducted on the raw Fisher-*Z* scores computed from each pair of endpoints. This ensured that effects were not driven by complex data manipulation or scaling and so the data were not adjusted to account for across session shifts in the similarity of all multivariate patterns (see Methods). Interestingly however, we did observe that Fisher-*Z* scores decreased from pre- to post-video across all pairs of endpoints regardless of condition, in each ROI (see figure at https://osf.io/2y3pm). This is reflected by a notable session effect in each mixed-effects model indicating reduced levels of similarity between non-associated endpoints (i.e. endpoints not belonging to the same location, video, or experimental condition); minimum effect size: *t2736* = −1.529, *p* = .126, *BF_01_* = 1.655, *d* = 0.319. As the size of this session effect was relatively large, the *Session*Overlap* and *Session*Overlap*Behaviour* interactions involved *less* of a decrease in similarity scores relative to all other conditions (see Figure 3).

Given that similarity scores decrease across all endpoint-pairs, it is unlikely that the session effect was a direct result of our video manipulation (i.e. learning induced neural differentiation). A mass differentiation on this scale would imply implausibly large amounts of information gain as the uniqueness (or entropy) of all neural representations would have to increase. Instead, it is more likely that the reduced levels of similarity were caused by systematically higher levels of noise in the second session. Most significantly, increases in temperature caused by RF absorption during scanning will shift the thermal equilibrium that governs how many hydrogen nuclei are aligned to the external magnetic field (*B_0_*), and can therefore contribute to the MR signal (see https://osf.io/8kns6/). In this case, we would expect to see similar shifts in the level of similarity across the entire brain. To test this, we measured pattern similarity in the Genu of the corpus callosum, a region that should exhibit negligible levels of BOLD activity. Based on a seed voxel at MNI = [0, 26, 6], multivariate patterns were taken from the 122 white matter voxels closest to that seed in native space. The size of this ROI was chosen to reflect the average size of our *a priori* regions of interest. A mixed-effects regression model of these data did indeed show reduced levels of neural similarity from session 1 to session 2; *t_2739_* = −2.167, *p* = .030, *BF_01_* = 0.651, *d* = −0.452 (similar in magnitude to the session effect in all other regions; see https://osf.io/p9qzx/).

In sum, we conclude that the overall decrease in pattern similarity across sessions was not driven by any meaningful change in neural representations and, once controlled for, reveals a significant increase in pattern similarity in both the right PHC and RSC in the overlap condition, indicative of viewpoint independent representations.

#### Laterality of RSA effects

The above analyses identified location-based representations in both right-hemisphere ROIs but no similar effects in the left hemisphere. Given this, we explored whether each RSA effect was significantly stronger in the right versus the left hemisphere. Comparing the *Session*Overlap* effects in the PHC did indeed reveal a significantly stronger effect in the right hemisphere (*t_5390.5_* = 3.798, *p* < .001, *BF_01_* = 0.028, *d* = 0.792). Similarly, comparing the *Session*Overlap*Behaviour* interactions in the RSC revealed a significantly stronger effect in the right hemisphere (*t_5427.4_* = 2.708, *p* = .007, *BF_01_* = 0.251, *d* = 0.565). Note, Robertson et al. (2016) collapsed their analyses across hemisphere, potentially masking laterality effects. These results are consistent with observations and theoretical models that the right hemisphere may preferentially process spatial information in humans as a consequence of predominantly left lateralised language function (Shulman et al., 2010; Smith & Milnek, 1981; Vallortigara & Rogers, 2005).

#### Searchlight RSA

The searchlight analysis that tested for a *Session*Overlap* interaction across the whole brain revealed one small cluster in the right inferior occipital gyrus (area V4; *t_21_* = 4.78, *p_FWE_* = .010, *k* = 38, *BF_01_* < 0.003, *d* = 0.997). However, when BOLD similarity in the cluster was modelled with the full mixed-effects analysis described above, the *Session*Overlap* effect was found to not be significant (*t_2740_* = 1.734, *p* = .083, uncorrected, *BF_01_* = 1.259, *d* = 0.361). Model parameter estimates suggested that the searchlight effect was driven by below baseline BOLD similarity in the overlap condition before the videos were shown (95% CI: [-0.116, −0.021]), a result that is not consistent with any effect of interest. No other areas showed a significant *Session*Overlap* or *Session*Overlap*Behaviour* interaction in the searchlight analysis. Nonetheless, both of the previously reported effects in the PHC and RSC are evident in the searchlight analysis at subthreshold levels (*t_21_* > 2, *d* > 0.417; see https://neurovault.org/collections/4819/).

### Univariate responses to endpoints

We investigated whether each of our ROIs produced univariate BOLD activations consistent with a *Session*Condition* interaction, or a 3-way interaction with behaviour. No such effects were found; all *F’s* < 1.140, *p’s* > .288. Furthermore, a mass univariate analysis testing for these effects at the whole brain level yielded no significant activations.

### Univariate responses to videos

Finally, we investigated whether univariate BOLD responses to the video clips differed between the overlap and no-overlap conditions, or as a function of scene memory in the post-scanner test. A group-level model was specified with predictors for: (1) video type (overlap vs no-overlap), (2) post-video performance in matching A1 and A2 endpoints, and (3) the interaction between video type and behavioural performance. This revealed two clusters that produced significantly greater BOLD responses during overlap vs no-overlap videos (Figure 5, hot colours). The largest of these peaked in the medial prefrontal cortex and extended into the anterior cingulate, left frontal pole, and left middle frontal gyrus (*t_21_* = 5.53, *p_FWE_* < .001, *k* = 600, *BF_01_* < 0.001, *d* = 1.153). The second cluster peaked in the left supramarginal gyrus (*t_21_* = 5.40, *p_FWE_* = .004, *k* = 185, *BF_01_* = 0.001, *d* = 1.126), adjacent to a smaller, sub-threshold effect in the left angular gyrus.

**Figure 5.**
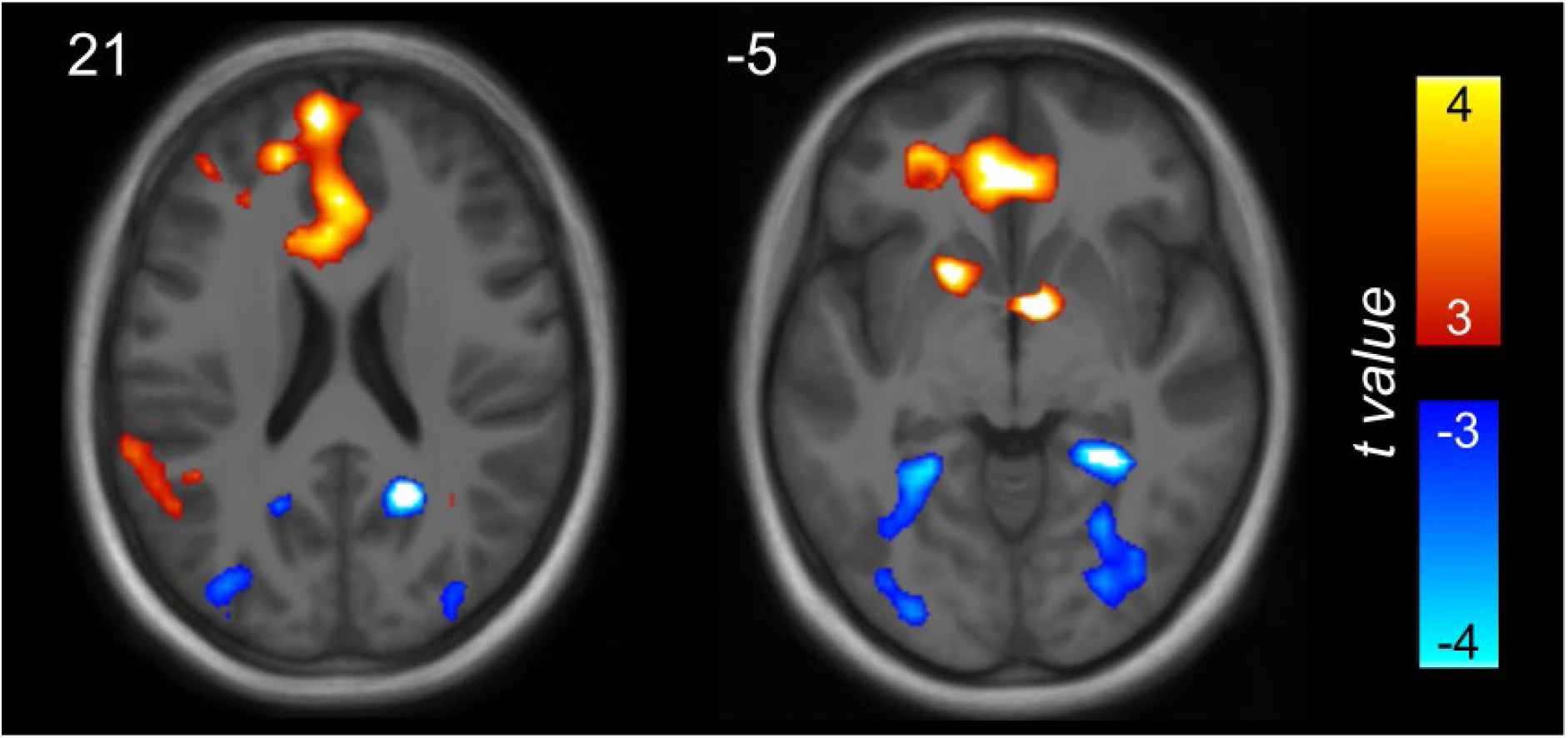
Univariate BOLD effects showing differences in activity between the two video conditions (thresholded at *t_21_* > 3, *p* < .004 uncorrected, *BF_01_* < 0.143, *d* > 0.626). Hot colours indicate areas showing a greater response to overlap vs no-overlap videos. Cool colours indicate areas showing a greater response to no-overlap vs overlap videos. An unthresholded statistical map of this contrast is available at https://neurovault.org/collections/4819.

No effects for the reverse contrast (i.e., no-overlap > overlap) reached statistical significance at the whole-brain level (subthreshold effects shown in Figure 5, cool colours). However, a small volume correction for the parahippocampal and retrosplenial cortices bilaterally revealed two clusters with a significant no-overlap > overlap effect. These were found in the right RSC (*t_21_* = −4.84, *p_FWE_* = .032, *k* = 26, *BF_01_* = 0.003, *d* = −1.001) and right PHC (*t_21_* = −4.77, *p_FWE_* = .026, *k* = 30, *BF_01_* = 0.003, *d* = −0.995), extending into the fusiform gyrus. Subthreshold effects for the no-overlap > overlap contrast were also evident in the left RSC and PHC. These results were mirrored by a linear mixed-effects model contrasting overlap and no-overlap video responses averaged across each ROI in native space. Here, both the right PHC and right RSC exhibited greater BOLD activity in the no-overlap video condition relative to the overlap condition (*t_42_* = −3.638, *p_FWE_* = .003, *BF_01_* = 0.039, *d* = −0.759 and *t_42_* = −3.499, *p_FWE_* = .004, *BF_01_* = 0.052, *d* = −0.730, respectively). Effects in the left PHC and left RSC were below threshold and considerably weaker (*t_42_* = −1.828, *p_FWE_* = .299, *BF_01_* = 1.101, *d* = −0.381 and *t_42_* = −2.212, *p_FWE_* = .130, *BF_01_* = 0.605, *d* = −0.461, respectively). Neither the whole-brain analysis nor the mixed-effects model identified BOLD responses to the videos that significantly correlated with memory performance in the post-scanner test.

In sum, we saw greater activity in the medial prefrontal cortex during the overlap videos relative to the no-overlap videos. In contrast, the PHC and RSC, showed greater activity during the no-overlap relative to overlap videos. In other words, the medial posterior regions that showed *increased* pattern similarity following presentation of the overlap video showed *decreased* activity while participants were watching the videos.

## Discussion

We show that scene-selective brain regions rapidly learn location-based representations of novel environments by integrating information across different viewpoints. Once participants observed the spatial relationship between two viewpoints from a given location, BOLD pattern similarity between viewpoints increased in right PHC and RSC, implying the emergence of location-based representations. In the right PHC, these representations appeared regardless of whether participants could identify which scenes were from the same location. In contrast, representations in the right RSC only emerged for scene pairs that participants could subsequently identify as being from the same location.

The results provide further evidence that PHC and RSC support spatial representations that are not solely driven by visual features in a scene (Marchette et al., 2015; Robertson et al., 2016; Vass & Epstein, 2013; c.f. Watson, Hartley, & Andrews, 2017). Using a similar panoramic video manipulation, Robertson et al. (2016) suggested that the RSC and OPA maintain viewpoint-independent representations, but found a more general associative effect in the PHC. Our results further identify the PHC in this process and highlight that RSC representations are more tightly linked to behaviour. Note, the OPA was not one of our *a priori* regions of interest, and we therefore make no claims in relation to this region supporting location-based representations (see Methods – Regions of Interest for further details). Our results also place constraints on models that describe how location-based representations are used. Unlike Robertson et al., we show that viewpoint-independent representations are evoked during passive viewing, in the absence of any explicit memory task, (though we cannot rule out the possibility that participants engaged in active imagery, as explicitly required in Robertson et al. see below).

Furthermore, we show that the learning of location-based representations can take place rapidly (in a single scanning session), with few exposures to the spatial layout of a location. Consistent with this, the firing fields of place cells have been shown to emerge rapidly in the rodent hippocampus (Monaco et al., 2014). Novel locations, where rats engaged in head-scanning behaviour (i.e., exploration), were associated with place fields the next time the rat visited the same location. Our results provide evidence that location-based representations form after only three learning exposures to the videos. Although we were specifically interested in the emergence of viewpoint-independent spatial representations, a similar approach could be used to track the emergence of viewpoint-independent representations of other stimulus categories (e.g., objects or faces; see Clarke et al., 2016 for a similar approach), opening the door to understanding how such representations are formed, or modulated, across the visual system.

We also found that right RSC only exhibited location-based representations when participants were able to identify which scenes belonged to that location in a post-scanner test (PHC representations emerged regardless of behavioural performance on the post-scanner test). This implies that the ability to identify differing scenes as from the same location is perhaps more dependent on representations in RSC than PHC. Computational models hold that medial posterior and temporal regions (including the PHC and RSC) perform distinct but complementary functions in support of spatial navigation and imagery (Bicanski & Burgess, 2018; Byrne et al., 2007). Specifically, PHC is thought to represent allocentric information related to the spatial geometry of the environment. Conversely, the posterior parietal cortex supports egocentric representations that allows the organism to actively navigate. The RSC transforms allocentric representations in the MTL into egocentric representations in the parietal cortex (and vice versa). Critically, the models predict that spatial navigation and planning is carried out in an egocentric reference frame. Thus, the RSC is critical to the translation of allocentric, to more behaviourally relevant, egocentric, information.

Our task required participants to match distinct scenes from the same location. This likely requires transformation from the presented egocentric viewpoint to an allocentric representation (ego-to-allocentric; i.e. A1 to the allocentric representation A*). In turn, the allocentric representation may allow for the retrieval of the associated viewpoint from the same location (allo-to-egocentric; i.e. A* to A2). Under this assumption, the RSC is likely to be more tightly coupled to behaviour relative to the PHC, as shown in the present data. This is because allocentric representations in the PHC only require the initial ego-to-allocentric transformation to be retrieved (A1-A*). If only the ego-to-allocentric transformation occurs, participants will not be able to perform the task. As such, it is possible to see evidence for allocentric PHC representations in the absence of accurate behaviour. For allocentric representations in the RSC to be retrieved, both the initial ego-to-allocentric (A1-A*), and subsequent allo-to-egocentric (A*-A2), transformation is required. If both transformations occur, then participants should be able to perform the task accurately. Thus, location-based representations in the RSC may only be seen in the presence of accurate behaviour and may reflect the transformation between reference frames rather than reflecting an allocentric representation per se.

A related possibility is that, during the passive viewing of specific scenes, participants engaged in active imagery of the associated scenes leading to subsequent improvements in behaviour for scenes from the same location. However, we note that the task did not explicitly require memory retrieval; participants responded to odd-ball targets leaving little time to for active imagery (see Linde-Domingo, Treder, Kerrén, & Wimber, 2019). Additionally, participants would only be able to engage in active imagery on the overlap trials alone. Despite this, we did not observe any univariate BOLD effects indicative of additional processing on these trials. As such, the activation of these representations does not appear to depend on any task-specific memory demands. It is possible that the retrieval of PHC representations (i.e., ego-to-allocentric mapping) occurs relatively automatically, consistent with the proposal that allocentric representations in the MTL are automatically updated during self-motion in an environment (Bicanski & Burgess, 2018; Byrne et al., 2007). However, the retrieval of associated egocentric representations (i.e., allo-to-egocentric mapping) may not occur automatically during passive viewing, consistent with the observation that viewpoint-independent representations in the RSC are abolished when participants engage in a task that prevents them from active retrieval of spatial information (Marchette et al., 2015). Importantly, both of the above accounts are consistent with the proposal that the RSC plays a critical role in mapping between allocentric and egocentric representations.

Although consistent with models of allocentric processing, it is possible that the location-based representations we observed reflect other forms of associative learning (e.g. O’Reilly & Rudy, 2001). On this view, scene A1 may become bound to A2 via a simple associative representation such that, after seeing the videos, A2 is covertly retrieved when presented with A1 (leading to increased pattern similarity). However, contrary to our findings, this simple account may also predict increased similarity in the no-overlap condition, where B1 and C2 are shown in the same video - particularly given that models of associative learning often rely on prediction error signals to account for incidental encoding (Den Ouden, Friston, Daw, McIntosh, & Stephan, 2009), which could be strongest in the no-overlap condition. A second possibility is that the overlapping content in the overlap videos (relative to the no-overlap videos) increases the probability of a direct association between A1 and A2. Indeed, it is the overlapping content that likely drives the increase in pattern similarity between overlap endpoints. Our current study is not able to discern whether the resulting “location-based” representations are associative, or truly allocentric, in nature.

In terms of associative learning, a related possibility is that the overlapping content supports a more complex transitive representation (e.g. A1-AX and A2-AX where X is the overlapping scene in the centre of the panorama). On this account, presentation of A1 cues the retrieval of AX and subsequently A2 (similar in nature to AB-AC inference paradigms; see Horner & Burgess, 2014; Joensen et al., 2019; Schlichting, Mumford, & Preston, 2015; Schlichting, Zeithamova, & Preston, 2014; Zeithamova, Dominick, & Preston, 2012). Representations that encode these transitive relationships between scenes are possible and may support spatial navigation but are not directly predicted by models of spatial memory (Bicanski & Burgess, 2018; Byrne et al., 2007). Further, the hippocampus and mPFC are more typically associated with transitive inference (Schlichting et al., 2015, 2014; Zeithamova et al., 2012), yet we only found evidence of location-based representations in scene-selective regions. Additionally, Robertson et al. have demonstrated that associative memory for scenes belonging to different locations is poor (comparable to their no-overlap condition) even when those scenes are presented in a ‘morphed’ panorama such that they are associated with a common context. As such, our data are suggestive of processes that go beyond associative or transitive learning and provide support for models of allocentric processing, although we cannot rule out an “associative” explanation.

Finally, it is noteworthy that certain non-spatial models may be able to account for our findings. In particular, models of directed attention may predict increased levels of pattern similarity in the overlap condition if the overlap videos alerted participants to visual features that are shared across scenes (e.g. Luo, Roads, & Love, 2020; Mack, Preston, & Love, 2013). Further work will be needed to fully establish the true nature of the location-based representations that we report here. To fully match all visual features across scenes in each condition, one possibility would be to experimentally manipulate the central section of continuous panoramas so that no coherent spatial representation can be learned. Furthermore, to fully distinguish between allocentric and transitive (A1-AX-A2) representations, an imaging study incorporating the panoramic morph manipulation used by Robertson et al. may be used.

While we directly link to computational models of spatial navigation and imagery, as well as rodent studies on spatial navigation, it is important to note that we have assessed pattern similarity during visual presentation of static scenes. This is a common approach in human fMRI (Bonner & Epstein, 2017; Marchette et al., 2015; Marchette et al., 2014; Robertson et al., 2016), as it allows one to control for many potential experimental confounds that might be present in a more ecologically valid experimental setting (e.g. using virtual reality; Doeller, Barry, & Burgess, 2010; Julian et al., 2016). However, this approach has the issue of being further removed from real-world spatial navigation. Interestingly, we saw evidence for increases in pattern similarity despite using a low-level attentional task, speaking to the potential automaticity of retrieving more location-based representations. Across the literature there are numerous examples of evidence for spatial learning in humans and rodents during goal directed navigation (Aoki, Igata, Ikegaya, & Sasaki, 2019; Howard et al., 2014), non-goal directed navigation (e.g. O’Keefe & Dostrovsky, 1971; Tolman, 1948), mental imagery (e.g. Bellmund, Deuker, Schro, & Doeller, 2016; Horner et al., 2016), and viewing of static images (e.g. Marchette et al., 2015; Robertson et al., 2016; Vass & Epstein, 2013). Our study adds to this growing literature suggesting that these representations can be assessed across dsiverse experimental environments with multiple methodologies.

The PHC has been proposed to represent several complementary spatial representations, including geometric information regarding one’s location in relation to bearings and distances to environmental features (e.g., boundaries; Park, Brady, Greene, & Oliva, 2011). The representations that we observed in PHC may reflect enriched spatial representations relating specific scenes to environmental features outside the current field-of-view. Also consistent with our results, PHC may represent spatial contexts more broadly (Epstein & Vass, 2014). The experimental manipulation used here could be modified to learn novel locations in the same spatial context, potentially dissociating between the above accounts. A further proposal is that viewpoint-independent representations in the PHC reflect prominent landmarks that are visible across viewpoints (Marchette et al., 2015). While this proposal yields similar predictions to above, it is less able to account for our finding of shared representations of views that did not contain any of the same landmarks.

Our PHC results are somewhat inconsistent with those of Robertson et al. (2016). Whereas our similarity increases were specific to the overlap condition, Robertson et al. saw effects in both their overlap and no-overlap conditions. One possibility is that our results reflect a Type II error, in that we failed to find an effect in the no-overlap condition when one is present. A second possibility is that Robertson et al. either (1) found an effect in the no-overlap condition when one is not present (i.e., a Type I error), or (2) failed to find a similarity effect in the overlap condition that was significantly larger than in the no-overlap condition (a Type II error). Notably, the ‘overlap > no-overlap’ effect size that we observed in the PHC is considerably larger than the same effect reported by Robertson et al. (0.610 relative to −0.062), and more in line with their RSC and OPA effects (0.470 and 0.415 respectively). Thus, it seems plausible that the disagreement stems from a Type I error in Robertson et al. Despite this, without further information, it is not possible to draw clear conclusions.

However, one important caveat is that we also saw evidence for a “same-location” effect in the PHC *before* learning had occurred. This effect was seen despite controlling for visual similarity across stimuli using the GIST descriptor, accounting for pixel-wise correlations in luminance and colour content, and despite participants being unable to identify which endpoints were from the same location prior to the videos. It is therefore possible that the PHC effects in Robertson et al. could have been driven by a similar effect not dependent on learning. This underlines the importance of including a pre- vs post-learning estimate of pattern similarity, to definitively rule out trivial effects driven by pre-existing similarities between images that are difficult to control for.

RSC representations may reflect the retrieval of spatial or conceptual information associated with the environment (Marchette et al., 2015). Further evidence suggests that the RSC contains multiple viewpoint dependent and independent (Vass & Epstein, 2013), as well as local and global (Jacob et al., 2017; Marchette et al., 2014), spatial representations. This multitude of representations fits with the proposed role of the RSC as a transformation circuit, mapping between allocentric and egocentric representations. The heterogeneity of representations, relative to the PHC, may also be a further reason why we did not see clear evidence for location-based representations without taking behaviour into account. Our RSC results are consistent with those of Robertson et al. in that they saw a clear effect following more extensive learning across two days (where behavioural performance was likely higher than in our study). However, we extend these findings to show that these effects are specifically associated with the locations that each individual participant has learned (i.e., a within-participant correlation that is consistent across participants). Regardless of the exact nature of such representations, our results provide clear evidence that we can track their emergence in both PHC and RSC.

Although more explorative, we also examined activity during learning of new spatial relationships (i.e., video presentation). BOLD activations in medial posterior brain regions (including but not limited to the PHC and RSC ROIs) were greater for no-overlap videos compared to overlap videos. This effect perhaps reflects greater fMRI adaptation during the overlap videos since they presented the central viewpoint of the panorama more frequently than no-overlap videos (Figure 1). However, it is interesting that the same cortical regions that showed *increased* pattern similarity following presentation of the overlap video showed *decreased* activity when participants were watching the videos. This underlines the complex relationship between univariate activity during learning, and resultant changes in patterns of activity following learning. More theoretically driven research would be needed to provide a robust explanation for this finding.

Additionally, we found that mPFC showed greater BOLD response in the overlap than no-overlap condition. This may reflect a mnemonic integration process that guides the learning of viewpoint-independent representations. Similar effects in mPFC have been observed in tasks that require integrating overlapping memories to support inference and generalisation (Milivojevic, Vicente-Grabovetsky, & Doeller, 2015; Schlichting et al., 2015). Indeed, mPFC has been implicated in detecting new information that is congruent with previously learnt material so that it can be integrated into a generalised representation (van Kesteren, Ruiter, Fernández, & Henson, 2012). Our results are broadly in line with this proposal, where mPFC may be detecting the presence of overlapping spatial information during the overlap videos, resulting in the integration of previously learnt representations into more coherent viewpoint-independent representations in posterior medial regions. Despite this, our results do not exclude the possibility that mPFC activations reflect disinhibition from medial-posterior inputs (which showed reduced activity), or attentional differences related to the behavioural task.

We have shown that brain regions in the scene network, specifically right PHC and RSC, rapidly learn representations of novel environments by integrating information across different viewpoints. They appear to be relatively viewpoint-independent in that they become active regardless of which part of an environment is in the current field-of-view. We show that PHC and RSC have potentially dissociable roles, consistent with models that propose the RSC plays a role in translating viewpoint-independent representations into a behavioural-relevant egocentric reference frame. Finally, our experimental approach allows for tracking the emergence of viewpoint-independent representations across the visual system.

## Acknowledgments

We thank all the staff at the York Neuroimaging Centre (YNiC) for their assistance in running this project. We are also grateful to Tim Andrews and Tom Hartley for early discussions regarding experimental design. AJH is funded by the Wellcome Trust (204277/Z/16/Z) and ESRC (ES/R007454/1). BHJ is funded a PhD studentship awarded by the Department of Psychology, University of York.

## References

Andersson, J. L. R., Hutton, C., Ashburner, J., Turner, R., & Friston, K. (2001). Modeling geometric deformations in EPI time series. NeuroImage, 13(5), 903–919. https://doi.org/10.1006/nimg.2001.0746

Aoki, Y., Igata, H., Ikegaya, Y., & Sasaki, T. (2019). The Integration of Goal-Directed Signals onto Spatial Maps of Hippocampal Place Cells Article The Integration of Goal-Directed Signals onto Spatial Maps of Hippocampal Place Cells. CellReports, 27(5), 1516–1527.e5. https://doi.org/10.1016/j.celrep.2019.04.002

Ashburner, J. (2007). A fast diffeomorphic image registration algorithm. NeuroImage, 38(1), 95–113. https://doi.org/10.1016/j.neuroimage.2007.07.007

Baayen, R. H., Davidson, D. J., & Bates, D. M. (2008). Mixed-effects modeling with crossed random effects for subjects and items. Journal of Memory and Language, 59(4), 390–412. https://doi.org/10.1016/j.jml.2007.12.005

Bellmund, J., Deuker, L., Schro, T. N., & Doeller, C. F. (2016). Grid-cell representations in mental simulation. (5:e17089), 1–21. https://doi.org/10.7554/eLife.17089

Bicanski, A., & Burgess, N. (2018). A neural-level model of spatial memory and imagery. ELife, (7). https://doi.org/10.7554/eLife.33752

Bonner, M. F., & Epstein, R. A. (2017). Coding of navigational affordances in the human visual system. Proceedings of the National Academy of Sciences, 114(18), 4793–4798. https://doi.org/10.1073/pnas.1618228114

Burgess, N., Becker, S., King, J. A., & O’Keefe, J. (2001). Memory for events and their spatial context: Models and experiments. Philosophical Transactions of the Royal Society B: Biological Sciences, 356(1413), 1493–1503. https://doi.org/10.1098/rstb.2001.0948

Byrne, P., Becker, S., & Burgess, N. (2007). Remembering the past and imagining the future: A neural model of spatial memory and imagery. Psychological Review, 114(2), 340–375. https://doi.org/10.1037/0033-295X.114.2.340

Calton, J. L., & Taube, J. S. (2009). Where am I and how will I get there from here? A role for posterior parietal cortex in the integration of spatial information and route planning. Neurobiology of Learning and Memory, 91(2), 186–196. https://doi.org/10.1016/j.nlm.2008.09.015

Clarke, A., Pell, P. J., Ranganath, C., & Tyler, L. K. (2016). Learning Warps Object Representations in the Ventral Temporal Cortex. Journal of Cognitive Neuroscience, 28(7), 1010–1023. https://doi.org/10.1162/jocn

Den Ouden, H. E. M., Friston, K. J., Daw, N. D., McIntosh, A. R., & Stephan, K. E. (2009). A dual role for prediction error in associative learning. Cerebral Cortex, 19(5), 1175–1185. https://doi.org/10.1093/cercor/bhn161

Doeller, C. F., Barry, C., & Burgess, N. (2010). Evidence for grid cells in a human memory network. Nature, 463(February). https://doi.org/10.1038/nature08704

Eichenbaum, H. (2004). Hippocampus: Cognitive processes and neural representations that underlie declarative memory. Neuron, 44(1), 109–120. https://doi.org/10.1016/j.neuron.2004.08.028

Epstein, R. A., Patai, E. Z., Julian, J. B., & Spiers, H. J. (2017). The cognitive map in humans: Spatial navigation and beyond. Nature Neuroscience, 20(11), 1504–1513. https://doi.org/10.1038/nn.4656

Epstein, R. A., & Vass, L. K. (2014). Neural systems for landmark-based wayfinding in humans. Philosophical Transactions of the Royal Society B: Biological Sciences, 369(1635). https://doi.org/10.1098/rstb.2012.0533

Gelman, A., Jakulin, A., Pittau, M. G., & Su, Y. S. (2008). A weakly informative default prior distribution for logistic and other regression models. Annals of Applied Statistics, 2(4), 1360–1383. https://doi.org/10.1214/08-AOAS191

Hannula, D. E., & Ranganath, C. (2009). The Eyes Have It: Hippocampal Activity Predicts Expression of Memory in Eye Movements. Neuron, 63(5), 592–599. https://doi.org/10.1016/j.neuron.2009.08.025

Henson, R. N., & Gagnepain, P. (2010). Predictive, interactive multiple memory systems. Hippocampus, 20(11), 1315–1326. https://doi.org/10.1002/hipo.20857

Horner, A. J., Bisby, J. A., Zotow, E., Bush, D., Burgess, N., Horner, A. J.,… Burgess, N. (2016). Grid-like Processing of Imagined Navigation Report Grid-like Processing of Imagined Navigation. Current Biology, 26(6), 842–847. https://doi.org/10.1016/j.cub.2016.01.042

Horner, A. J., & Burgess, N. (2014). Report Pattern Completion in Multielement Event Engrams. Current Biology, 24(9), 988–992. https://doi.org/10.1016/j.cub.2014.03.012

Howard, L. R., Javadi, A. H., Yu, Y., Mill, R. D., Morrison, L. C., Knight, R.,… Spiers, H. J. (2014). The Hippocampus and Entorhinal Cortex Encode the Path and Euclidean Distances to Goals during Navigation. 1331–1340. https://doi.org/10.1016/j.cub.2014.05.001

Hutton, C., Bork, A., Josephs, O., Deichmann, R., Ashburner, J., & Turner, R. (2002). Image distortion correction in fMRI: A quantitative evaluation. NeuroImage, 16(1), 217–240. https://doi.org/10.1006/nimg.2001.1054

Jacob, P. Y., Casali, G., Spieser, L., Page, H., Overington, D., & Jeffery, K. (2017). An independent, landmark-dominated head-direction signal in dysgranular retrosplenial cortex. Nature Neuroscience, 20(2), 173–175. https://doi.org/10.1038/nn.4465

Joensen, B. H., Gaskell, M. G., Horner, A. J., Joensen, B. H., Gaskell, M. G., & Horner, A. J. (2019). Journal of Experimental Psychology: General Episodic Events United We Fall: All-or-None Forgetting of Complex Episodic Events.

Julian, J. B., Fedorenko, E., Webster, J., & Kanwisher, N. (2012). An algorithmic method for functionally defining regions of interest in the ventral visual pathway. NeuroImage, 60(4), 2357–2364. https://doi.org/10.1016/j.neuroimage.2012.02.055

Julian, Joshua B., Keinath, A. T., Marchette, S. A., & Epstein, R. A. (2018). The Neurocognitive Basis of Spatial Reorientation. Current Biology, 23(17), R1059–R1073. https://doi.org/10.1016/j.cub.2018.04.057

Julian, Joshua B., Ryan, J., Hamilton, R. H., & Epstein, R. A. (2016). The Occipital Place Area Is Causally Involved in Representing Environmental Boundaries during Navigation. Current Biology, 26(8), 1104–1109. https://doi.org/10.1016/j.cub.2016.02.066

Kass, R. E., & Raftery, A. E. (1995). Bayes Factors Bayes Factors. Journal of the American Statistical Association ISSN:, 90(430), 773–795.

Kumaran, D., Hassabis, D., Spiers, H. J., Vann, S. D., Vargha-Khadem, F., & Maguire, E. A. (2007). Impaired spatial and non-spatial configural learning in patients with hippocampal pathology. Neuropsychologia, 45(12), 2699–2711. https://doi.org/10.1016/j.neuropsychologia.2007.04.007

Linde-Domingo, J., Treder, M. S., Kerrén, C., & Wimber, M. (2019). Evidence that neural information flow is reversed between object perception and object reconstruction from memory. Nature Communications, 10(1). https://doi.org/10.1038/s41467-018-08080-2

Luo, X., Roads, B. D., & Love, B. C. (2020). The Costs and Benefits of Goal-Directed Attention in Deep Convolutional Neural Networks. ArXiv. Retrieved from http://arxiv.org/abs/2002.02342

Mack, M. L., Preston, A. R., & Love, B. C. (2013). Decoding the brain’s algorithm for categorization from its neural implementation. Current Biology, 28(20), 2023–2027. https://doi.org/10.1016/j.cub.2013.08.035

Malcolm, G. L., Silson, E. H., Henry, J. R., & Baker, C. I. (2018). Transcranial Magnetic Stimulation to the Occipital Place Area Biases Gaze During Scene Viewing. Frontiers in Human Neuroscience, 12(May), 1–13. https://doi.org/10.3389/fnhum.2018.00189

Marchette, S. A., Vass, L. K., Ryan, J., & Epstein, R. A. (2015). Outside Looking In: Landmark Generalization in the Human Navigational System. Journal of Neuroscience, 35(44), 14896–14908. https://doi.org/10.1523/JNEUROSCI.2270-15.2015

Marchette, Steven A., Ryan, J., & Epstein, R. A. (2017). Schematic representations of local environmental space guide goal-directed navigation. Cognition, 158, 68–80. https://doi.org/10.1016/j.cognition.2016.10.005

Marchette, Steven A., Vass, L. K., Ryan, J., & Epstein, R. A. (2014). Anchoring the neural compass: Coding of local spatial reference frames in human medial parietal lobe. Nature Neuroscience, 17(11), 1598–1606. https://doi.org/10.1038/nn.3834

Milivojevic, B., Vicente-Grabovetsky, A., & Doeller, C. F. (2015). Insight reconfigures hippocampal-prefrontal memories. Current Biology, 25(7), 821–830. https://doi.org/10.1016/j.cub.2015.01.033

Monaco, J. D., Rao, G., Roth, E. D., & Knierim, J. J. (2014). Attentive scanning behavior drives one-trial potentiation of hippocampal place fields. Nature Neuroscience, 17(5), 725–731. https://doi.org/10.1038/nn.3687

Motley, S. E., Grossman, Y. S., Janssen, W. G. M., Baxter, M. G., Rapp, P. R., Dumitriu, D., & Morrison, J. H. (2018). Selective Loss of Thin Spines in Area 7a of the Primate Intraparietal Sulcus Predicts Age-Related Working Memory Impairment. The Journal of Neuroscience, 38(49), 10467–10478. https://doi.org/10.1523/jneurosci.1234-18.2018

O’Keefe, J, & Dostrovsky, J. (1971). The hippocampus as a spatial map: Preliminary evidence from unit activity in the freely-moving rat. Brain Research, (34), 171–175.

O’Keefe, John, & Burgess, N. (2005). Dual phase and rate coding in hippocampal place cells: Theoretical significance and relationship to entorhinal grid cells. Hippocampus, 15(7), 853–866. https://doi.org/10.1002/hipo.20115

O’Reilly, R., & Rudy, J. (2001). Conjunctive representations in Learnig and memory. Psychological Review, 108(2), 311–345. Retrieved from http://www.ncbi.nlm.nih.gov/pubmed/11381832

Oliva, A., & Torralba, A. (2001). Modeling the shape of the scene: A holistic representation of the spatial envelope. International Journal of Computer Vision, 42(3), 145–175. https://doi.org/10.1023/A:1011139631724

Park, S., Brady, T. F., Greene, M. R., & Oliva, A. (2011). Disentangling Scene Content from Spatial Boundary: Complementary Roles for the Parahippocampal Place Area and Lateral Occipital Complex in Representing Real-World Scenes. Journal of Neuroscience, 31(4), 1333–1340. https://doi.org/10.1523/JNEUROSCI.3885-10.2011

Robertson, C. E., Hermann, K. L., Mynick, A., Kravitz, D. J., & Kanwisher, N. (2016). Neural Representations Integrate the Current Field of View with the Remembered 360° Panorama in Scene-Selective Cortex. Current Biology, 26(18), 2463–2468. https://doi.org/10.1016/j.cub.2016.07.002

Schlichting, M. L., Mumford, J. A., & Preston, A. R. (2015). Learning-related representational changes reveal dissociable integration and separation signatures in the hippocampus and prefrontal cortex. Nature Communications, 6, 1–10. https://doi.org/10.1038/ncomms9151

Schlichting, M. L., Zeithamova, D., & Preston, A. R. (2014). CA 1 Subfield Contributions to Memory Integration and Inference. 1260, 1248–1260. https://doi.org/10.1002/hipo.22310

Shulman, G. L., Pope, D. L. W., Astafiev, S. V., McAvoy, M. P., Snyder, A. Z., & Corbetta, M. (2010). Right Hemisphere Dominance during Spatial Selective Attention and Target Detection Occurs Outside the Dorsal Frontoparietal Network. Journal of Neuroscience, 30(10), 3640–3651. https://doi.org/10.1523/jneurosci.4085-09.2010

Silson, E., Steel, A., & Baker, C. (2016). Scene selectivity and retinotopy in medial parietal cortex. Journal of Vision, 16(12), 528. https://doi.org/10.1167/16.12.528

Smith, M. Lou, & Milnek, B. (1981). The role of the right hippocampus spatial location in the recall of spatial location. Neuropsychologia, 19(6), 781–793.

Tolman, E. C. (1948). Cognitive Maps in Rats and Men. Psychological Review, 55(4), 189.

Tzourio-Mazoyer, N., Landeau, B., Papathanassiou, D., Crivello, F., Etard, O., Delcroix, N.,… Joliot, M. (2002). Automated anatomical labeling of activations in SPM using a macroscopic anatomical parcellation of the MNI MRI single-subject brain. NeuroImage, 15(1), 273–289. https://doi.org/10.1006/nimg.2001.0978

Vallortigara, G., & Rogers, L. (2005). Survival With an Asymmetrical Brain: Advantages and Disadvantages of Cerebral Lateralization. Behavioral and Brain Sciences, 28(4), 575–589. https://doi.org/10.1017/s0140525x05000105

Van Kesteren, M. T. R., Ruiter, D. J., Fernández, G., & Henson, R. N. (2012). How schema and novelty augment memory formation. Trends in Neurosciences, 35(4), 211–219. https://doi.org/10.1016/j.tins.2012.02.001

Vass, L. K., & Epstein, R. A. (2013). Abstract Representations of Location and Facing Direction in the Human Brain. Journal of Neuroscience, 33(14), 6133–6142. https://doi.org/10.1523/JNEUROSCI.3873-12.2013

Watson, D. M., Hartley, T., & Andrews, T. J. (2017). Patterns of response to scrambled scenes reveal the importance of visual properties in the organization of scene-selective cortex. Cortex, 92, 162–174. https://doi.org/10.1016/j.cortex.2017.04.011

Welch, B. L. (1947). The Generalization of “Student’s” Problem when Several Different Population Variances are Involved. Biometrika, 34(1), 28–35. https://doi.org/10.1093/biomet/34.1-2.28

Zeithamova, D., Dominick, A., & Preston, A. R. (2012). Hippocampal and Ventral Medial Prefrontal Activation during Retrieval-Mediated Learning Supports Novel Inference. 75, 168–179. https://doi.org/10.1016/j.neuron.2012.05.010

